# The anti-leprosy drug clofazimine reduces polyQ toxicity through activation of PPAR*γ*

**DOI:** 10.1101/2023.02.06.527298

**Authors:** Xuexin Li, Ivo Hernandez, Maria Häggblad, Louise Lidemalm, Lars Brautigam, Jose J. Lucas, Jordi Carreras-Puigvert, Daniela Hühn, Oscar Fernandez-Capetillo

## Abstract

PolyQ diseases are autosomal dominant neurodegenerative disorders caused by the expansion of CAG repeats. While of slow progression, these diseases are ultimately fatal and lack effective therapies. Here, we present our results from a High-Throughput chemical screen oriented to find drugs that lower the toxicity of a protein containing the first exon from the Huntington’s disease protein huntingtin (HTT) harboring 94 glutamines (Htt-Q_94_). Our screening identified the anti-leprosy drug clofazimine as a hit, which was subsequently validated in several in vitro models as well as in a zebrafish model of polyQ toxicity. Computational analyses of transcriptional signatures, together with molecular modeling and biochemical assays revealed that clofazimine is an agonist of the peroxisome proliferator activated receptor gamma (PPAR*γ*), previously suggested as a potential therapy for HD by stimulating mitochondrial biogenesis. Accordingly, clofazimine rescued the mitochondrial dysfunction triggered by Htt-Q_94_ expression. Together, our results support the potential of clofazimine repurposing for the treatment of PolyQ diseases.

## INTRODUCTION

Polyglutamine (polyQ) diseases include 9 inherited hereditary neurodegenerative syndromes that are caused by the expansion of Q-coding repeats within the exons of several seemingly unrelated genes [1]. One of these pathologies is Huntington’s disease (HD), being one of the most frequent neurodegenerative diseases with an incidence of 3-5 cases per 100.000 worldwide [2]. In HD, the disease is linked to the expansion of a CAG repeat within the first exon of huntingtin (HTT), which becomes pathogenic above 35 repeats with the severity of the disease correlating with repeat length [3, 4]. While HTT dysfunction has been proposed to contribute to HD [5, 6], an alternative hypothesis is that the pathology is caused by gain-of-function toxicity of the polyQ-bearing mutant HTT (mHTT). Accordingly, early studies showed that transgenic mice expressing a fragment of the exon 1 from mHTT including the expanded polyQ track suffered from motor dysfunction and premature death [7, 8]. Importantly, seminal work revealed that ectopic expression of polyQ expansions inserted in *HPRT*, a gene not mutated in patients, also led to neurodegeneration and premature death, highlighting the causal role of polyQ toxicity [9].

In what regards to the mechanisms of polyQ toxicity, this remains to be fully understood. An important feature of these expansions is their propensity to form insoluble aggregates that form intraneuronal inclusions, which were found in mouse models and also patients from several polyQ diseases including HD [10-12]. However, whether these inclusions are the real cause of the pathology has been the subject of intense debate, and it is clear that mHTT can also be toxic independently of the formation of large aggregates (reviewed in [1]). Regardless of whether it forms inclusions, mHTT has been shown to drive multiple cellular alterations in aspects such as mRNA transcription [13-15], protein degradation and post-translational modifications [16], synaptic function and plasticity [17-21] and mitochondrial activity [22-26].

Unfortunately, these mechanistic discoveries have not yet led to clinical improvements in the treatment of HD. The only approved treatments for HD, tetrabenazine and deutetrabenazine, are directed to alleviate the involuntary movements (chorea), but do not cure the disease [27, 28]. In this context, it becomes urgent to try to find novel therapies for polyQ diseases, an area of intense research. Efforts are spread among strategies trying to prevent mHTT aggregates or promote their clearance, as well as to targeting their downstream pathological effects (reviewed in [29]). Noteworthy, several of the unbiased chemical screens have been focused on the identification of compounds that lower polyQ aggregates in biochemical assays, which often lead to compounds that show toxicity per se when evaluated in in vivo models [30, 31]. Here, we present our results from a High-Throughput Imaging based drug-repurposing screening oriented to find compounds that reduce the toxicity of polyQ expansions.

## RESULTS

### A chemical screen for modulators of polyQ-toxicity

To conduct a chemical screen, we first generated an inducible system enabling the expression of an EGFP fusion protein containing the first exon of human HTT with an expanded polyQ tract of 94 glutamines (Htt-Q_94_ hereafter). The cDNA was cloned in a Tet-On gene expression system enabling the expression of Htt-Q_94_ upon the addition of doxycycline (dox). As polyQ expression is toxic for any cell type, the system was stably integrated in human osteosarcoma U2OS cells that are widely used in large chemical screens, and a clone with stringent regulated expression selected for further experiments (U2OS^Q94^). Before conducting the screen, we verified the Dox-inducible expression of Htt-Q_94_ as seen by a widespread accumulation of EGFP-expressing cells (**Fig. 1A**). Moreover, and as previously reported in similar setups, a 1-week treatment with dox led to the appearance of cells with Q_94_ aggregates (**Fig. 1A**, inset). At this time, Htt-Q_94_ expression led to a significant reduction in cell numbers, as quantified by detecting nuclei by High-Throughput Microscopy (**Fig. 1B**), confirming the toxicity driven by polyQ expression in this cell system.

**Figure 1.**
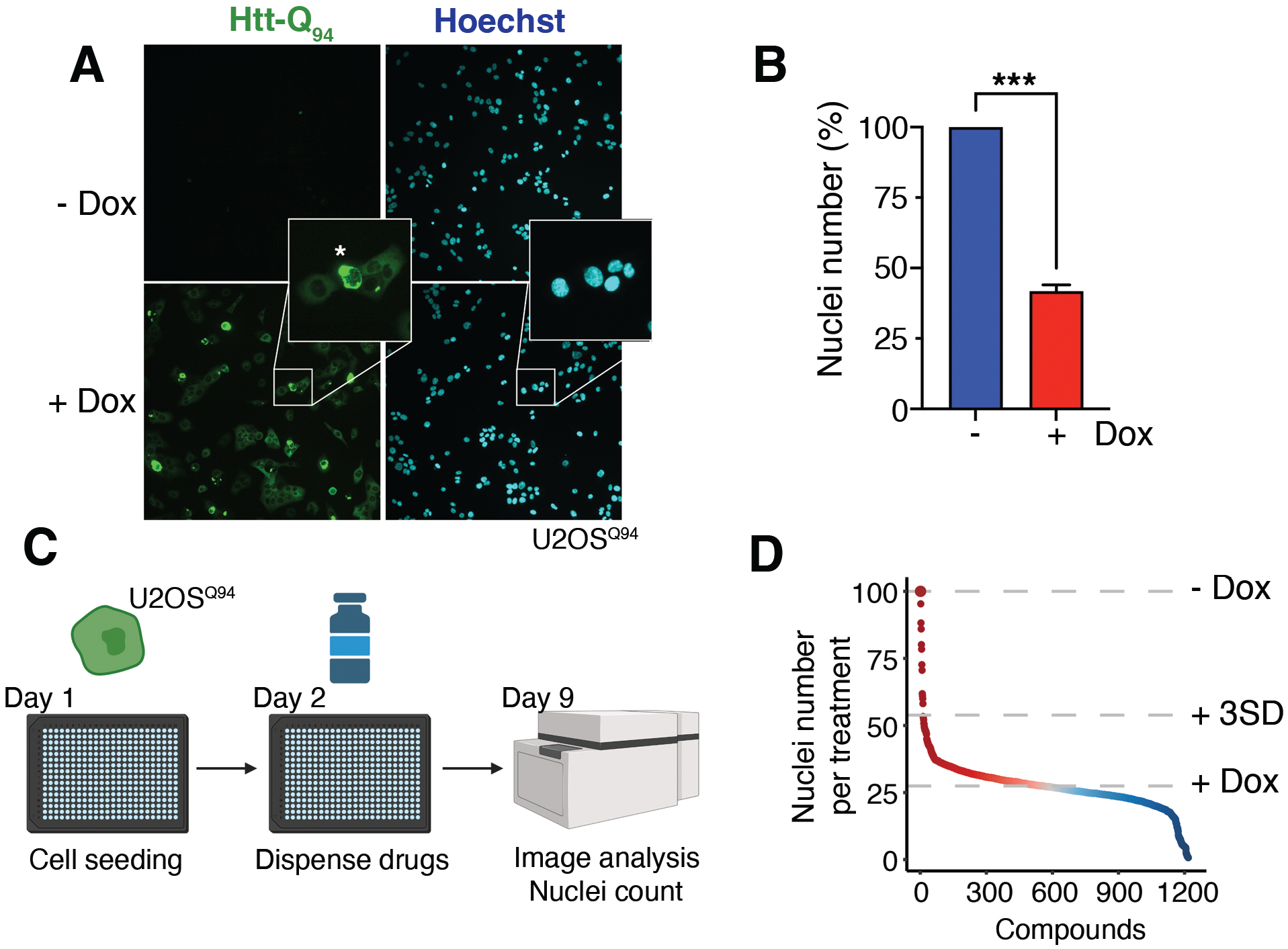
A chemical screen to identify modifiers of polyQ toxicity. (**A**) Representative image of Htt-Q_94_ expression (monitored by EGFP, green) in U2OS^Q94^ cells treated or not with dox (50 ng/ml) for 8 days. An inset illustrates the appearance of cells harbouring perinuclear aggregates Htt-Q_94_ (asterisk). Hoechst (blue) was used to stain DNA and enable the quantification of nuclei. (**B**) High-Throughput Microscopy (HTM)-mediated quantification of nuclei numbers from the data presented in (**A**). \*\*\**p*<0.001, t-test. (**C**) Pipeline of the chemical screen. On day 0, U2OS^Q94^ cells were seeded on 384-well plates. On the following day, cells were treated with dox (50 ng/ml) and with the compounds from the library at 1*μ*M. Nuclei numbers were quantified by HTM on day 9. Scattered controls of cells not treated with dox, or only treated with dox but without additional compounds were used for normalization. (**B**) Hit distribution of the screen described in (**C**). Compounds that led to an increase in nuclei numbers higher than 3SD when compared to the numbers founds on wells only treated with dox were taken for secondary validation (**Fig S1**).

The chemical library used combined 1,200 FDA-approved compounds and 94 additional drugs targeting components of the epigenetic machinery (**Table S1**). We added epigenetic drugs given that several of them have been found to be of potential for the treatment of HD and other neurodegenerative diseases [32]. To conduct the screen, U2OS^Q94^ cells were seeded on 384 well plates at 100 cells/well, and treated with dox (50 ng/ml) and the library compounds (1*μ*M) for 8 days. At this point, cells were fixed and DNA was stained with Hoechst enabling the quantification of nuclei (**Fig. 1D**). 35 compounds leading to an increase in nuclei numbers bigger that 3 SD from those found in the control wells (only treated with dox), were taken for a dose-response validation screen conducted at 0.5, 1, 5 and 10 *μ*M. From this secondary screen, 4 compounds showed a significant rescue of toxicity in at least 2 of the doses tested: promethazine (PRM), amodiaquine (AMD), clofazimine (CFZ) and troglitazone (TZD) (**Fig. S1A**).

### Clofazimine and troglitazone rescue polyQ-toxicity in vitro

Next, and to evaluate whether the compounds were able to present a sustained effect in reducing the toxicity associated to Htt-Q_94_ expression, we conducted clonogenic survival assays. These experiments confirmed that all 4 compounds increased the number of colonies in dox-treated U2OS^Q94^ cells (**Fig. 2A** and **Fig. S2**). Before entering into mechanistic analyses, we first wanted to discard hits that were acting by preventing the dox-dependent expression of Htt-Q_94_, an issue that we have previously faced when conducting similar screens using Tet-On systems [33]. To do so, we measured their effects on dox-induced Htt-Q_94_ levels both by immunofluorescence (IF) and western blotting (WB). While PRM and AMD significantly limited Htt-Q_94_ expression, CFZ and TZD rescued polyQ toxicity despite not affecting Htt-Q_94_ expression or the presence of its aggregates (**Fig. 2B, C**). We thus focused on these two compounds for subsequent analyses.

**Figure 2.**
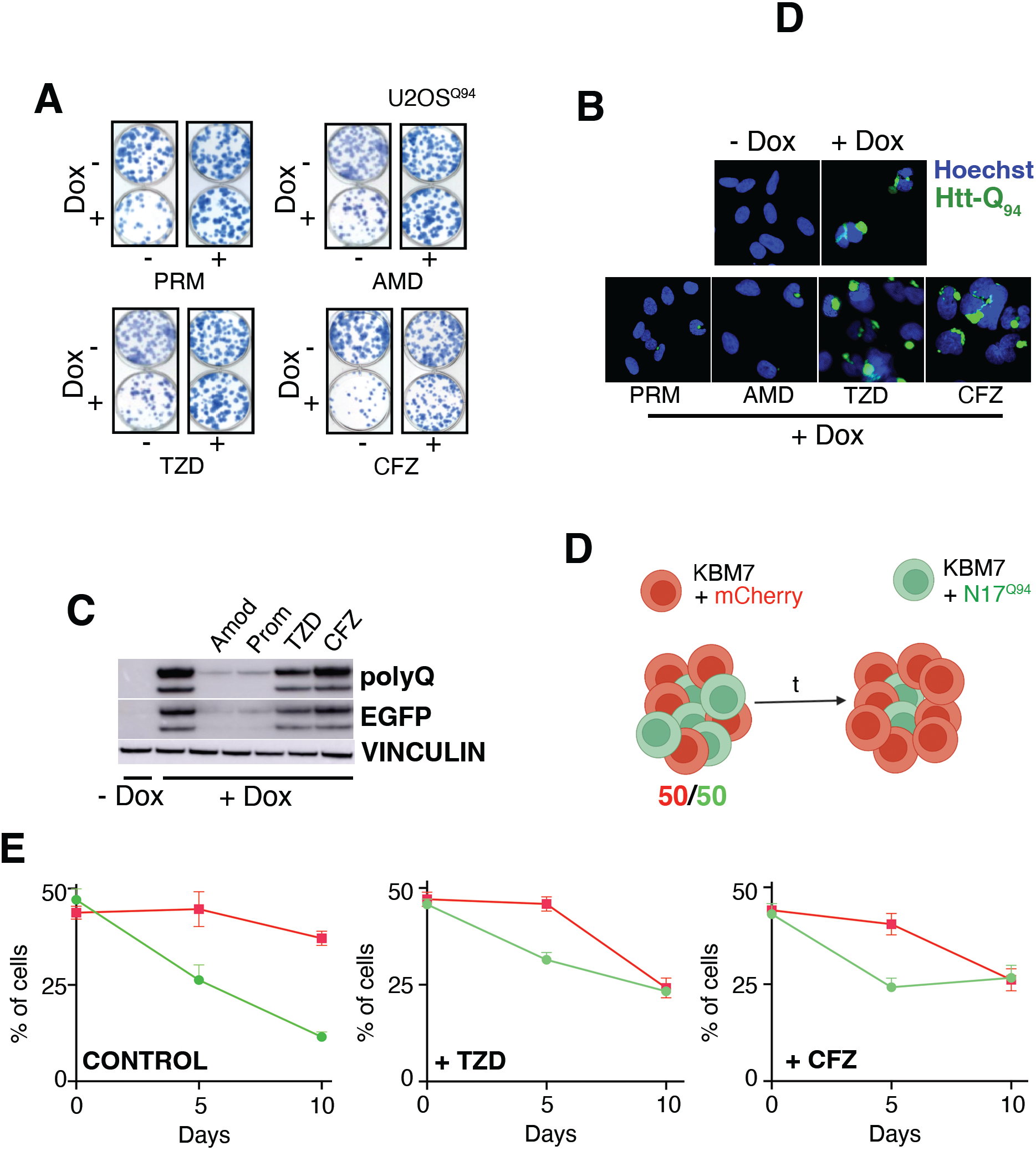
Clofazimine and troglitazone alleviate polyQ toxicity in vitro. (**A**) Representative images from clonogenic survival assays performed in U2OS^Q94^ cells, treated or not with dox (50ng/ml) and the indicated drugs at 2*μ*M for 12 days. The full dose-response dataset from the clonogenic assays is available at (**Fig. S2**). (**B**) Representative images of Htt-Q_94_ expression (detected by the EGFP signal, green) in U2OS^Q94^ cells, treated or not with dox (50ng/ml) and the indicated drugs at 5*μ*M for 8 days. Nuclei were stained with Hoechst (blue). (**C**) WB analysis of Htt-Q_94_ expression levels, monitored both with an anti-EGFP antibody or an antibody against polyQ peptides, in the experiment defined in (**B**). Vinculin levels were assessed as a loading control. (**D**) Scheme of the competition assay using KBM7 cells expressing either mCherry (red, control) or a fusion protein between Htt-Q_94_ and EGFP (green). When co-cultured, the percentage of Htt-Q_94_ progressively declines. (**E**) Data from the KBM7 competition experiment defined in (**D**), in cultures treated with DMSO (control), TZD or CFZ (at 5 *μ*M).

To further validate these results in vitro in an orthogonal model, we performed growth competition assays in the human leukemic KBM7 cell line. To do so, we co-cultured KBM7 cells expressing either mCherry or EGFP-Htt-Q_94_ (KBM7^Q94^) for 10 days. In the absence of drugs, the percentage of KBM7^Q94^ cells progressively declined, confirming that Htt-Q_94_ expression also impairs cellular fitness in this model. In contrast, treatment with CFZ or TZD rescued the relative decline of KBM7^Q94^ cells, confirming the in vitro effects of both these drugs in rescuing polyQ toxicity (**Fig. 2D**,**E**). Interestingly, one of these two compounds, TZD, is a well-established agonist of the peroxisome proliferator activated receptor gamma (PPAR*γ*), an approach that has been previously studied as a potential therapy for various neurodegenerative diseases including HD, confirming the usefulness of our screen to identify potential therapies [34-37, 26]. In contrast, CFZ, an antibiotic originally developed as a treatment for leprosy active against a wide range of mycobacteria [38], has not been previously investigated in the context neurodegeneration. We thus selected CFZ for further analyses.

### Clofazimine rescues polyQ-induced mitochondrial damage

To understand how CFZ treatment was rescuing polyQ toxicity we conducted transcriptomic analyses by RNA sequencing (RNAseq) in dox-induced U2OS^Q94^ cells treated or not with CFZ for 8 days. Interestingly, these analyses revealed a general impact of CFZ in boosting the expression of multiple factors related to mitochondria such as voltage dependent ion channels, translocases, subunits of the ATP synthase and components of mitochondrial translation (**Fig. 3A**). Consistently, Gene Set Enrichment Analyses revealed that CFZ treatment led to a significant enrichment of multiple pathways related to mitochondrial function (**Fig. 3B**) in dox-induced U2OS^Q94^ cells.

**Figure 3.**
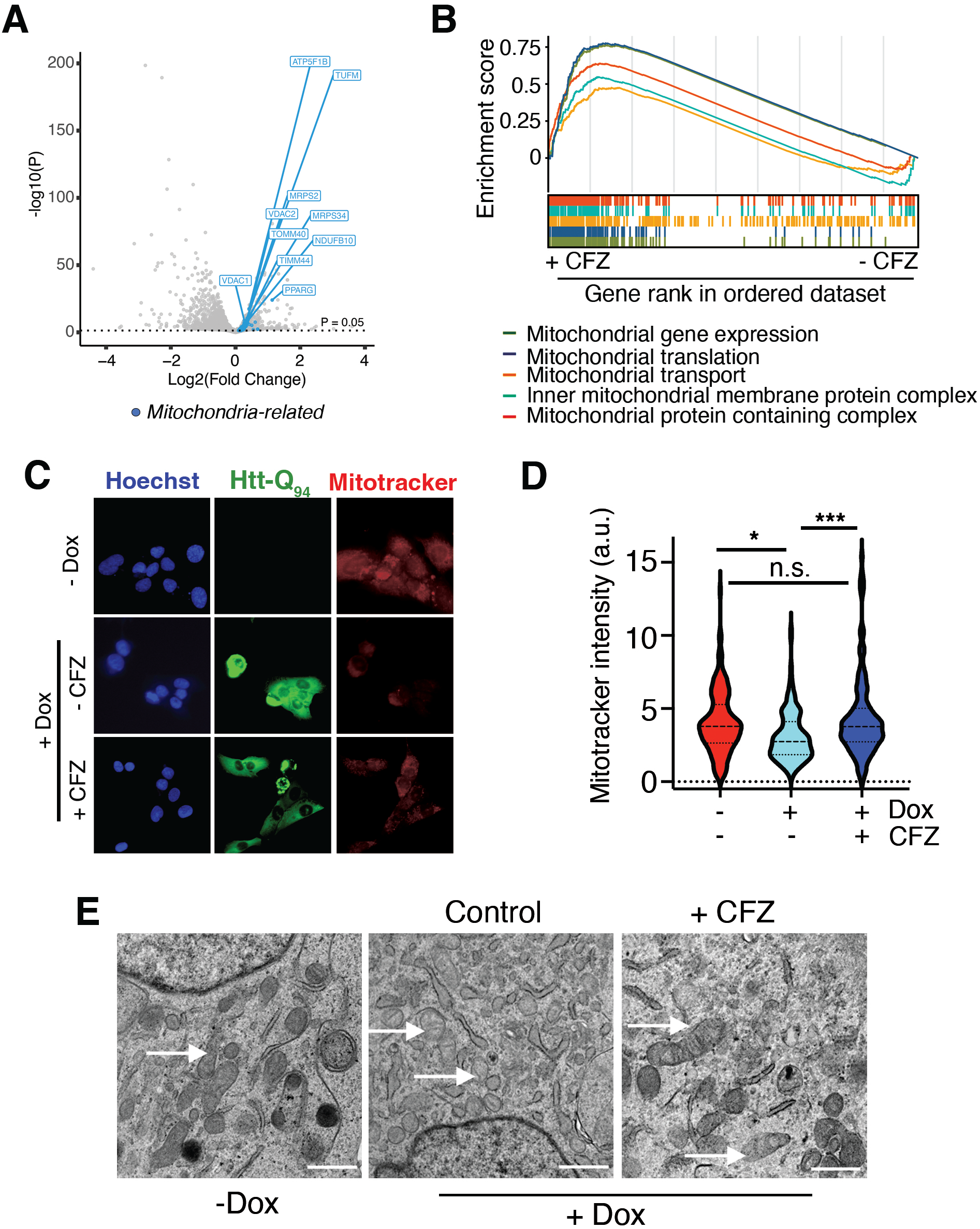
Clofazimine restores mitochondrial function in polyQ-expressing cells. (**A**) Volcano plot representing RNAseq data illustrating the impact of CFZ treatment (5 *μ*M; 8 days) in dox-induced U2OS^Q94^ cells. Genes above dotted line are differentially regulated (p < 0.05). Blue dots highlight mitochondria-related genes. (**B**) GSEA analyses from the experiment defined in A, illustrating the overall increase in mitochondria-related pathways upon CFZ treatment in dox-induced U2OS^Q94^ cells. (**C**) Representative images of Htt-Q_94_ expression (detected by the EGFP signal, green) or the mitotracker signal (red) in the of U2OS^Q94^ cells treated or not with dox (50 ng/ml) and CFZ (5 *μ*M). Nuclei were stained with Hoechst (blue). (**D**) HTM-dependent quantification of the cytoplasmic mitotracker signal per cell from the experiment defined in (**C**). \**p*<0.05, \*\*\**p*<0.001, t-test. (**E**) Representative images from transmission electron microscopy of U2OS^Q94^ cells treated or not with dox (50 ng/ml) and CFZ (5 *μ*M). Arrows indicate mitochondria, which are significantly altered upon expression, and improved upon a concomitant treatment with CFZ. Scale bar (white), represents 0,5 *μ*m.

To evaluate mitochondrial activity, we used mitotracker, a red dye that stains mitochondria in a membrane-potential-dependent manner [39]. In agreement with the mitochondrial dysfunction that has been repeatedly documented in cells from HD patients (reviewed in [40]), dox treatment led to a notable reduction of the mitotracker signal in U2OS^Q94^ cells, which was rescued by CFZ (**Fig. 3C**,**D**). Similarly, transmission electron microscopy analyses revealed that Htt-Q_94_ expression had a profound impact on the mitochondria of U2OS^Q94^ cells, characterized by swelling and substantial abnormalities in external membranes and cristae, all of which were rescued by CFZ (**Fig. 3E**).

### Clofazimine is a PPAR*γ* agonist

As mentioned, CFZ has been used as an antimycobacterial since the 1950s [41]. In addition, recent screens also identified that CFZ prevented infection by a wide range of viruses, including SARS-CoV-2 [42]. Surprisingly, despite its interesting medical properties, its target and mechanism of action remain unknown. To address this, we used the transcriptional signature of CFZ-treated U2OS cells to interrogate the Connectivity Map (CMap), a database from the Broad Institute at MIT that stores the transcriptional signatures of more than 5,000 drugs [43], aiming to identify drugs with a similar transcriptional impact. Interestingly, these analyses revealed an enrichment of PPAR*γ* agonists among the compounds presenting a transcriptional signature that resembled that of CFZ (**Fig. 4A**). In fact, PPAR*γ* itself was transcriptionally induced by CFZ in our transcriptomic analyses (**Fig. 3A**).

**Figure 4.**
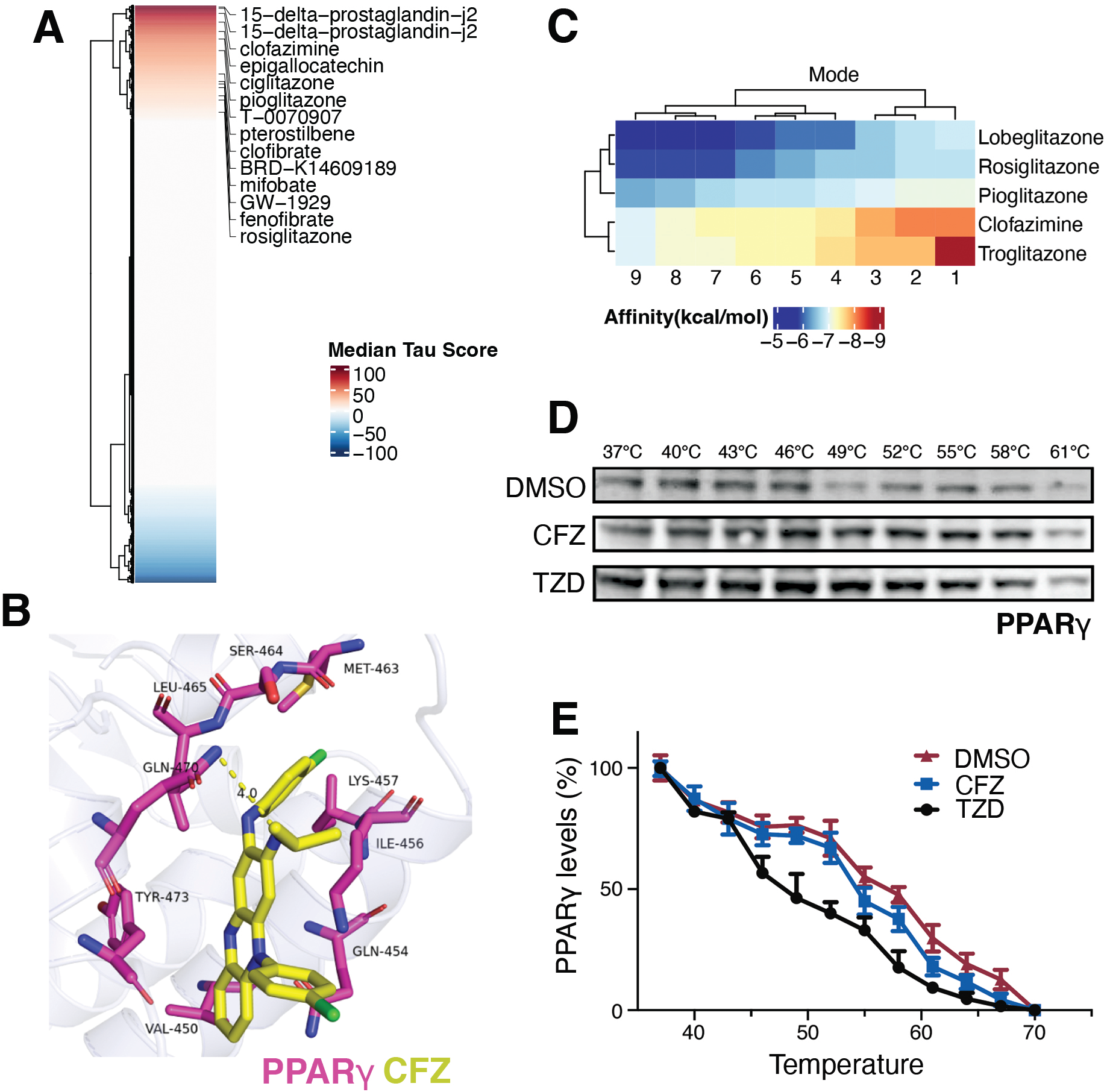
Identification of clofazimine as a PPAR*γ* agonist. (**A**) The transcriptional signature of CFZ-treated cells, was used as input to search for drugs exerting a similar transcriptional signature at the Connectivity Map database from the Broad Institute at MIT [43]. The panel indicates an enrichment of PPAR*γ* agonists among the drugs showing a transcriptional signature resembling that of CFZ. (**B**) Molecular docking illustrating the fitting of CFZ (yellow) in an allosteric pocket of PPAR*γ* (red). The interaction occurs through hydrophobic forces and the formation of a hydrogen bond with gln-470 (length of the bond, 4.0 Å). CFZ has hydrophobic interactions with tyr-473, val-450, gln-454, ile-456, lys-457, met-463, ser-464 and leu-465. (**C**) Binding affinities of CFZ and several PPARg agonists towards PPAR*γ*, based on the molecular docking experiment shown in (**B**). (**D**) Cellular thermal shift assay (CETSA) measuring the effects of TZD and CFZ on PPAR*γ* levels at increasing temperatures. Both compounds increased the thermal stability of PPAR*γ* when compared to the DMSO control. (**E**) Quantification of the CETSA studies shown in (**D**).

Consistent with bioinformatic analyses, molecular docking revealed that CFZ is able to bind to the same pocket in PPAR*γ* as other agonists such as TZ, and with a similar binding affinity (**Fig. 4B**,**C**). Furthermore, cellular thermal shift assays (CETSA) [44] indicated that TZ and CFZ were able to have a similar impact in stabilizing PPAR*γ* at increasing temperatures, supporting a direct interaction (**Fig 4D**,**E**). Together, these results demonstrate that CFZ can bind to PPAR*γ* and stimulate its activity.

### Clofazimine rescues polyQ toxicity in neurons and zebrafish

To further document the effect of CFZ in a neuronal model, we used SH-SY5Y neuroblastoma cells, which can be differentiated into a neuronal-like phenotype with retinoic acid (RA) [45]. Parental cells were infected with pLVX-UbC-rtTA-Htt-Q94-CFP lentiviruses, enabling dox-dependent expression of Htt-Q94-CFP (SH-SY5Y^Q94^). After 5 days of differentiation with 10 *μ*m RA, SH-SY5Y^Q94^ cells were treated with dox for 3 additional days. As in all previous models, Htt-Q_94_ expression had a profound impact on SH-SY5Y^Q94^ cells, exemplified by decreased cell numbers and a reduction in the mitotracker signal; all of which were rescued by treatment with CFZ (**Fig. 5A**,**B** and **Fig. S3**). Of note, CFZ had a significant effect in increasing cell numbers and mitochondrial activity also in SH-SY5Y^Q94^ cells that were not previously exposed to Dox, highlighting its potentially beneficial effects in other pathologies associated to neuronal dysfunction [46].

**Figure 5.**
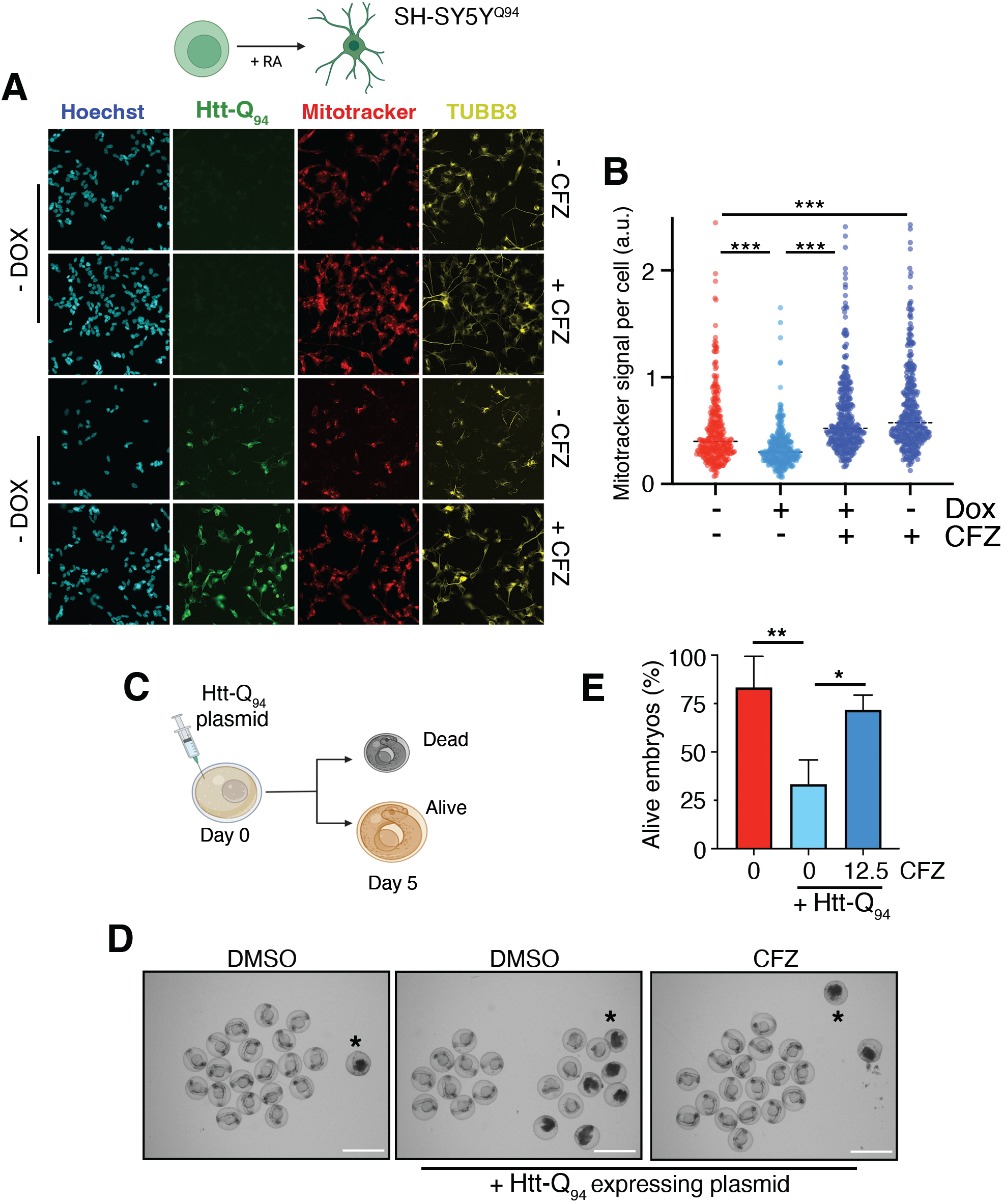
Clofazimine rescues polyQ toxicity in neurons and zebrafish. (**A**) Representative images of SH-SY5Y^Q94^ cells differentiated with RA (10*μ*M, 5 days), and subsequently treated with dox (35ng/ml) with or without CFZ (1*μ*M) for 3 additional days. Levels of Htt-Q_94_ (measured by the CFP signal), TUBB3 (yellow) and mitotracker (red) are shown. Hoechst (blue) was used to stain DNA and detect nuclei. An image of the entire well for this dataset, as well as the quantification of cell numbers is shown in **Fig. S3**. (**B**) HTM-dependent quantification of the cytoplasmic mitotracker signal per cell from the experiment defined in (**A**). (**C**) Scheme illustrating the pipeline followed to evaluate Htt-Q_94_ in developing zebrafish. Viability was monitored 5 days after microinjection with an Htt-Q_94_-CFP expressing plasmid. (**D**) Representative images of zebrafish embryos 5 days after the microinjection of the Htt-Q_94_-CFP expressing plasmid or DMSO. Note the accumulation of dead embryos (black asterisk) upon Htt-Q_94_-CFP expression, which was significantly rescued by CFZ (12.5 *μ*M). Scalebar (white) represents 1 mm. (**E**) Quantification from the experiment defined in (**C-D**). \**p*<0,05, \*\**p*<0.01, t-test.

Finally, we tested the impact of CFZ in alleviating polyQ toxicity in zebrafish. To this end, we used a previously developed plasmid enabling the expression of Htt-Q_94_-CFP [47]. On day 0, fertilized zebrafish eggs were injected with the plasmid and exposed to CFZ at 12,5 *μ*M (**Fig. 5C**). Consistent with previous studies [48, 49], transgenic Htt-Q_94_ expression led to substantial embryonic lethality in developing fish. Importantly, CFZ was able to significantly increase embryonic survival in Htt-Q_94_-transgenic embryos, confirming its effects in alleviating polyQ toxicity in vivo (**Fig. 5D**,**E**).

## DISCUSSION

As mentioned in the introduction, and despite the substantial advances made in understanding the molecular basis of polyQ-diseases, this has not yet led to effective treatments. Among others, substantial efforts are being dedicated to find therapeutic strategies to either reduce the expression of polyQ-containing proteins (e.g. antisense oligonucleotides (ASOs) or RNA interference), or that aim to either prevent the formation of polyQ aggregates or promote their clearance (reviewed in [29]). Our approach was rather to identify molecules capable of reducing the toxicity of polyQ-bearing proteins. In this regard, a similar approach was conducted by the Taylor laboratory where they searched for molecules that reduced apoptosis triggered by the expression of a truncated androgen receptor containing a 112-glutamine repeat in HEK 293T cells [50]. In our screen model, U2OS, Htt-Q_94_ did not trigger apoptosis but rather cell cycle arrest. Interestingly, we observed that the severity of this phenotype more acute when cells were sed at low densities, perhaps reflecting that the formation of polyQ aggregates is also enhanced at sub-confluence [51].

The usefulness of our approach was supported by the fact that we were able to identify compounds previously known to modulate the severity of polyQ pathology in preclinical models such as TZD [34, 35, 37, 26]. Unfortunately, and despite being originally approved for the treatment of diabetes, TZD was later removed from the marked due to hepatic toxicity [52]. Nevertheless, cumulative data supporting that activation of the PPAR*γ*/PDC1a axis is a fruitful therapeutic approach for the treatment of neurodegenerative diseases [46], emphasizes the need of discovering new PPAR*γ* agonists that hopefully overcome the initial toxicities. In this regard, our work indicates that CFZ is a PPAR*γ* agonist, with a similar binding affinity as TZD, but which is seemingly safe as it is in clinical use for the treatment of infectious diseases. Of note, one limitation of CFZ is its poor efficacy in crossing the blood-brain barrier (BBB), which has limited its efficacy for the treatment of infections in the central nervous system. In this regard, there are already efforts dedicated to circumvent this problem such as nanoparticle-based formulations of CFZ [53]. In any case, our work suggests that CFZ could be a useful alternative to TZD for the treatment of pathologies outside the CNS. In summary, our study further indicates the potential of PPAR*γ* stimulation to reduce the severity of pathologies of polyQ-diseases, and that these effects are primarily related to restoring mitochondrial function. In addition, our work adds a new example of the possibilities offered by drug repurposing to identify medically approved drugs that could be investigated in the context of neurodegenerative diseases. While acknowledging the current pharmacological limitations of the drug, we believe that exploring the efficacy of CFZ or its derivatives in polyQ diseases deserves further preclinical work.

## MATERIAL AND METHODS

### Cell culture, transfection and chemicals

All cells were grown at 37°C in a humidified air atmosphere with 5% CO_2_. U2OS (human osteosarcoma) cell line was cultured in DMEM +Glutamax (Thermo Fisher Scientific), 10% FBS and 1 % Penicillin/Streptomycin. U2OS^Q94^ cells were cultured in DMEM +Glutamax supplemented with 10% Tet system approved FBS (Takara, 631368) and 1% Penicillin/Streptomycin, selected with zeocin and Blasticidin S. KBM7^Q94^ and KBM7-mCherry cells were cultured in IMDM (Thermo Fisher Scientific), supplemented with 10% FCS and 1 % Penicillin/Streptomycin. SH-SY5Y^Q94^ cells were cultured in DMEM/F-12 (Thermo Fisher Scientific), 10% Tet system approved FBS and 1 % Penicillin/Streptomycin. For the transfection of U2OS cells, Lipofectamine2000 transfection reagent (Thermo Fisher Scientific) was used following standard protocol. For the transfection of KBM7 cells, Amaxa Nucleofector kit (Reactive L, X-001 program) was used, and 5×10^5^ cells with 10mg of indicated plasmids were transfected according to the manufacturer’s protocol. For the lentiviral transduction of SH-SY5Y cells, pLVX-UbC-rtTA-Htt-Q94-CFP vector was co-transfected in HEK293T cells using Lipofectamine2000 with packaging vectors pMD2.G (Addgene, #12259) and psPAX2 (Addgene, #12260). Lentiviral supernatants were collected 36 hours after transfection, filtered and immediately used for transduction.

### Plasmids

Human Htt-exon1-Q94 fragment from pTreTight-Htt94Q-CFP (Addgene, #23966) was cloned into pBlueScript SK+ (kind gift of Eva Brinkman) by using HindIII and BamHI sites to generate an intermediate plasmid, pBlueScript SK-polyQ94. PINTO-polyQ94-GFP was cloned by pBlueScript SK-polyQ94 and pINTO-N-GFP using KpnI and NotI sites. pcDNA3.1-EGFP-poly94 was cloned by using pcDNA3.1-mCherry (Addgene, #128744) [54] and pINTO-polyQ94-GFP with AflII and NotI sites. To clone plasmid for SH-SY5Y cell infection, polyQ94-CFP was cloned into pBlueScript SK+ by using XbaI and EcoRI sites to generate an intermediate plasmid SK-polyQ94-CFP. pLVX-UbC-rtTA-polyQ94-CFP was cloned by SK-polyQ94-CFP and pLVX-UbC-rtTA-Ngn2:2A:Ascl1 (Addgene, #127289) using the dual NotI sites. The genetic construct of zebrafish plasmid is based on the vector pDEST-Tol2-PA2-CMV-AB-mCh (Addgene, #160435) [55] in which the Abeta peptide was exchanged to EGFP-polyQ94 from pcDNA3.1-EGFP-polyQ94. Lentiviral packaging vectors pMD2.G (Addgene, #12259) and psPAX2 (Addgene, #12260) were used.

### High-throughput Screening (HTS)

Plate and liquid handling were performed using Echo550 (Labcyte, USA), Viaflo 384 (Integra Biosciences, Japan), Multiflo FX Multi-Mode Dispenser (BioTek, USA), and Hydrospeed washer (Tecan, Switzerland). Cells were seeded in black 384-well plates with clear bottom (BD Falcon, #353962). Compound libraries were provided by the Chemical Biology Consortium Sweden (CBCS). The chemical collection used in the primary screening contained 1,122 medically approved compounds from the Prestwick library and 94 epigenetic-drugs available at CBCS collections (the list of compounds is available at **Table S1**). For the primary screen, U2OS^Q94^ cells were trypsinized and resuspended in culture medium. The cell suspension (100 cells in 30 μl/well) was dispensed into 384-well plates and exposed to a final concentration of 1μM of compounds diluted in dimethyl sulfoxide (DMSO) for 8 days. Cells were with 4% PFA and nuclei were stained with 2 μM Hoechst 33342 for 15 min in the dark.

Plates were imaged using an IN Cell Analyzer 2200 system (GE Healthcare, USA) with a 10× objective, 4 images per well were acquired, covering the whole well. Images were analyzed with the open-source software CellProfiler [56] using a custom-made pipeline for the detection of nulei count and cytoplasmic GFP signal. The analysis considered the integrated intensity of cytoplasmic GFP staining and the nuclei count. All values were normalized to DMSO conditions within each plate. Then, the mean value for each compound in triplicates was calculated, representing a single measurement per compound. For the validation screen, U2OS^Q94^ cells were exposed to 4 concentrations, 0.5, 1, 3, and 10 μM, of the selected hits for 8 days. The validation was conducted in triplicates and images were analyzed as described above. Statistical analysis of imagining data was conducted using Graphpad Prism software.

### Immunoblotting

Cell pellets were lysed in in RIPA buffer (Thermo Fisher Scientific) supplemented with protease and phosphatase inhibitor cocktail (Roche), sonicated for 5 min and centrifuged at 4°C, 14000 rpm for 15 min. 30μg whole-cell extracts were separated by SDS–PAGE and transferred onto Nitrocellulose membrane (Bio-Rad). After blocking in 5% milk in TBS-T, indicated antibodies were diluted in blocking buffer and incubated overnight at 4°C. The following dilutions of primary antibodies was used: GFP (1:300, Abcam, #ab290), Polyglutamine (1:1000, Sigma-Aldrich, #P1874), PPAR-γ (1:250, Abcam, a#b45036), Vinculin (1:2000, Abcam, #ab130007). The signal associated to HRP-conjugated secondary antibodies (ThermoFisher, mouse #31430 and rabbit #31460) was developed with a SuperSignal West Pico PLUS Chemiluminescent Substrate kit (ThermoFisher, #34580), and analyzed in an Amersham Imager 600.

### Flow cytometry

For the analysis of the competition assay of KBM7 _Q94_ and KBM7-mCherry in, 4x 10^4^ KBM7 _Q94_ and KBM7-mCherry cells were mixed at a 1:1 ratio and were seeded in T175 flasks. Cells were treated with CFZ and TZD at indicated concentrations. After 5 or 15 days, cells were analysed for green or red fluorescence by flow cytometry (Bio-Rad S3e cell sorter). Data were processed with the Flow Jo 10 software to calculate the percentage of each cell population.

### Viability assays

For clonogenic survival assays, 500 cells per well were plated in 6-well tissue culture plates in the corresponding culture medium. Cells were treated with the indicated concentrations of drugs and maintained with the compounds for 12 days, changing the medium every 3-4 days, and then fixed and stained with 0.4% methylene blue in methanol for 30 min.

### RNA-seq and data analysis

Total RNA was extracted from cell pellets using a Purelink RNA Mini Kit (Invitrogen #12183025) following manufacturer’s instructions. Total RNA was subjected to quality control with Agilent Tapestation (#G2991BA). To construct libraries suitable for Illumina sequencing, an Illumina stranded mRNA prep ligation sample preparation protocol was used with an starting concentration of total RNA between 25-1000 ng. The protocol includes mRNA isolation, cDNA synthesis, ligation of adapters and amplification of indexed libraries. The yield and quality of the amplified libraries was analyzed using a Qubit by Thermo Fisher and the quality of the library was checked using the Agilent Tapestation. Indexed cDNA libraries were normalized and combined, and pools were sequenced using an Illumina platform. STAR [57] was used for sequence alignment based on the GRCh38 DNA primary assembly reference build [58], and quantification was done using featureCounts [59] with reference build GRCh38.101 [58].

Differential expression (DE) analyses between the groups were performed using DESeq2 [60]. Generalized linear model (GLM) was fitted to the expression data and shrunken log2fold-change (LFC) using adaptive Student’s t prior shrinkage estimator [60, 61]. Multiple testing correction was done using Benjamini-Hochberg (BH) method [62]. GSEA analysis was performed on the gene-level statistics from the DE analyses results against the molecular signatures from the Molecular Signatures Database (MSigDB) v7.5.1 [63, 64] and the Reactome database [65]. Specifically, signatures from the ontology gene set C5 of MSigDB, containing Gene Ontology (GO)-derived gene sets [66, 67], as well as the complete gene sets from the Reactome database, retrieved from fgsea 1.20.0 [68], were used. GSEA analysis was carried out using clusterProfiler 4.2.2 [69] to identify enriched terms. The transcriptional signatures, identified to be the sets of top up/down-regulated 150 genes (BH p-adjusted values < 0.05, ranked by LFC) from the DE analysis outcomes, were separately used as inputs to the Connectivity Map (CMap) Query clue.io tool (https://clue.io/) [70] to identify drugs with signatures similar to that clofazimine.

### Molecular docking

The 3D crystal structure of PPAR-γ was downloaded from the Protein Data Bank (http://www.rcsb.com; #3ET0). Autodock vina was used to removed water molecules and add missing side or back chains and residues. Chemical structures were downloaded from the ZINC database of molecular structures for virtual screens (www.zinc15.org). Molecular-docking was conducted by Auto Dock vina and the binding energy of each ligand was analyzed based on binding free energies and root mean square deviation (RMSD) values. Top nine binding energies of each ligand were listed.

### Zebrafish study

To test the impact of CFZ in alleviating polyQ toxicity in zebrafish. On day0, the injection mix (25ng/ul transposase, 50ng/ul vector, 0.3% phenol red) was injected into 150 eggs. The compounds were added in to the E3 medium at 12,5 *μ*M, CFZ treatment naïve injected group and DMSO treated un-injected group are also inclouded. After 24h, dead embryos/well are counted and imaged. The experiment was triplicated.

### Statistics

Statistical parameters and tests are reported in the Figures and corresponding Figure Legends. Statistical analysis was done using GraphPad Prism version 8.0 (GraphPad Software Inc). One-way-ANOVA was performed for all the datasets that required comparison among multiple data points within a given experimental condition.

### Data availability

RNA sequencing data associated to this work are accessible at the GEO repository, under accession number GSE222758.

## ACKNOWLEDGEMENTS

We thank the Chemical Biology Consortium Sweden for their help with the chemical libraries. Research was funded by grants from the Cancerfonden foundation (CAN 2018/381) and the Swedish Research Council (VR) (538-2014-31) to OF; and the Spanish Ministry of Science and Innovation and Spanish Ministry of Science (PID2021-123141OB-I00 MCIU/AEI/FEDER, UE) to JJL.

## AUTHOR CONTRIBUTIONS

X.L. contributed to most experiments and data analyses and to the preparation of the figures. I.H. and J.J.L. helped with animal models. M.H. helped with the chemical screen. L.L. provided technical help in several of the in vitro experiments. L.L. helped with zebrafish experiments. J.C.-P. contributed to the original design of the screen. D.H. contributed to most of the analyses and supervision of the project. O.F-C. supervised the study and wrote the MS.

