## Supplementary material for "The anti-leprosy drug clofazimine reduces polyQ toxicity through activation of PPAR*γ*": Figures S1-3

**Figure S1**. **Dose-response evaluation of the primary hits.** 35 hits identified in our primary screen (described in **Fig. 1**) as potentially increasing nuclei numbers in dox-treated U2OS$^{Q94}$ cells, were evaluated using the same pipeline at increasing doses (0, 0.5, 1, 5 and 10 $\mu$M). Nuclei numbers were quantified at day 8 by HTM. Data show that four of the initial hits (highlighted by a red arrow) presented a dose-response increase in nuclei numbers in dox-treated U2OS$^{Q94}$ cells, and were selected for further analyses.

Li *et al.* **Fig S1**

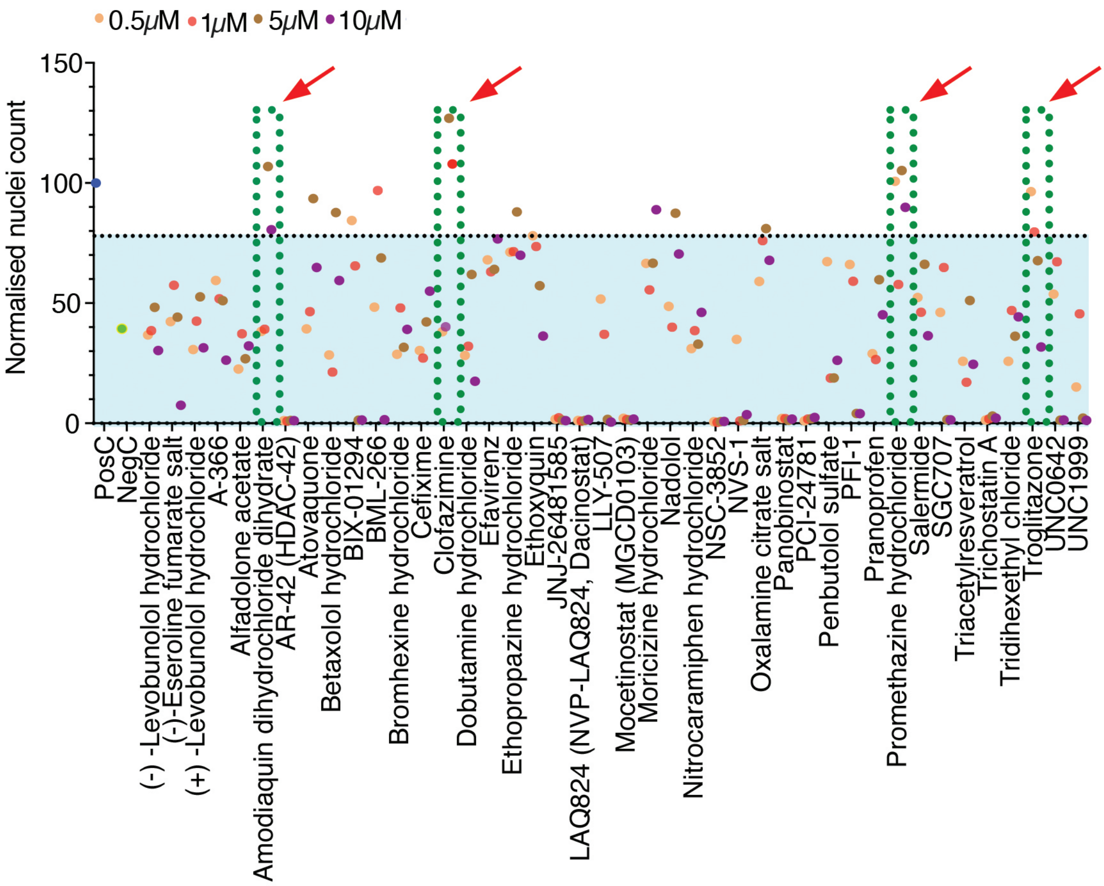

**Figure S2**. **Clonogenic assays of the 4 initial hits.** Images from the full dose-response clonogenic survival assays performed in U2OS$^{Q94}$ cells, treated or not with dox (50ng/ml) and the indicated drugs for 12 days. The 2 μM dose was shown as a representative example in **Fig. 2A**.

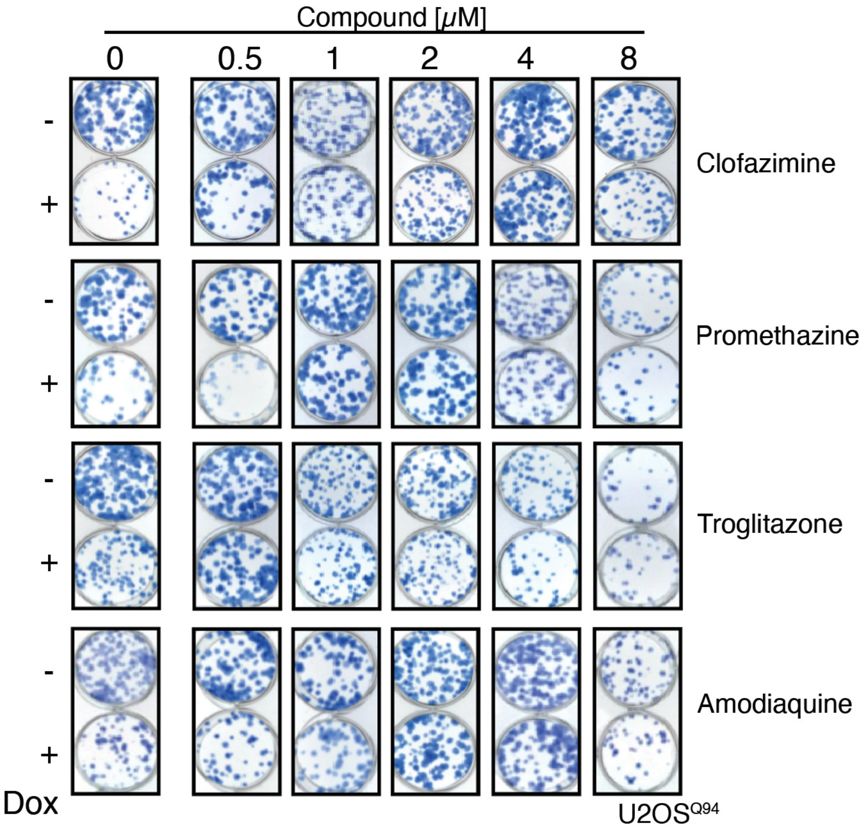
Compound [μM]
0
0.5
1
2
4
8
-
+
Clofazimine
Promethazine
Troglitazone
Amodiaquine
Dox
U2OS^Q94

**Figure S3**. **Rescue of polyQ toxicity by CFZ in SH-SY5Y$^{Q94}$ cells.** Representative images from the entire well of a 384-well plate of SH-SY5Y$^{Q94}$ cells differentiated with RA (10μM, 5 days), and subsequently treated with dox (35ng/ml) with or without CFZ (1μM) for 3 additional days. Levels of Htt-$Q_{94}$ (measured by the CFP signal), TUBB3 (yellow) and mitotracker (red) are shown. Hoechst was used to stain DNA and visualize nuclei. Magnified insets from this dataset are shown in **Fig. 5A**. (**B**) HTM-dependent quantification of the number of nuclei per well from the experiment defined in (**A**).

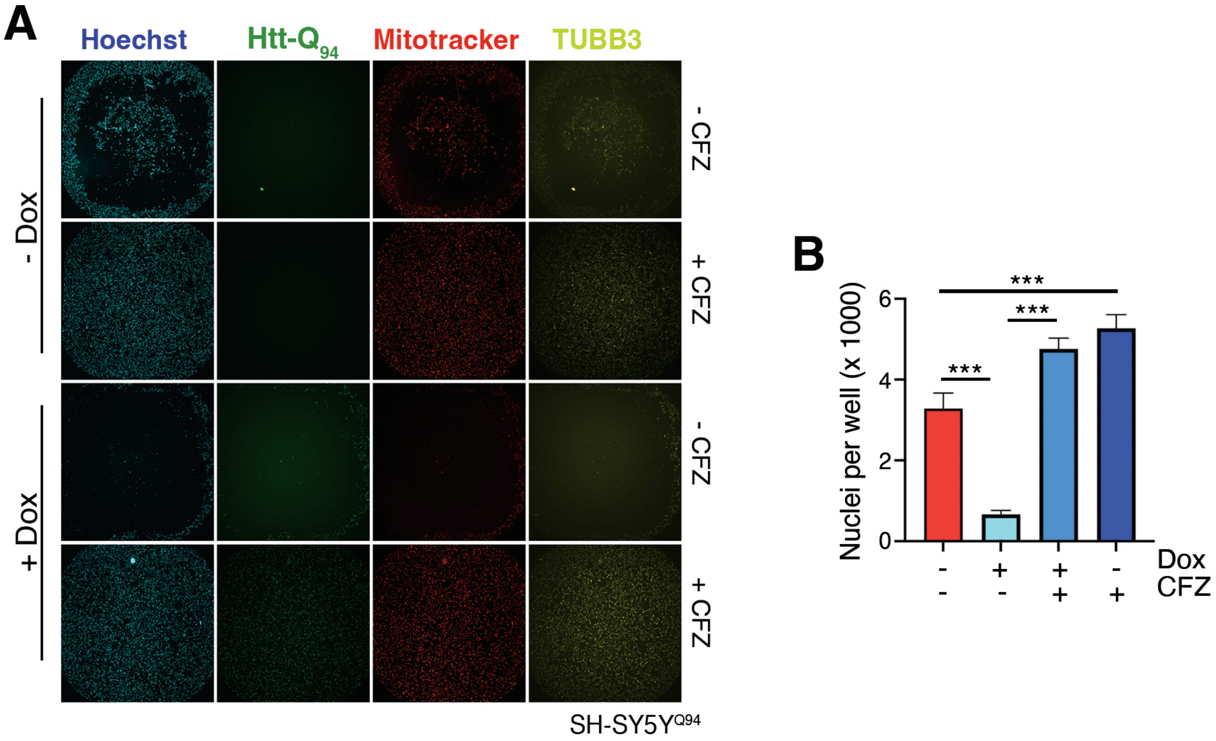
A
Hoechst
Htt-Q94
Mitotracker
TUBB3
- Dox
+ Dox
- CFZ
+ CFZ
- CFZ
+ CFZ
SH-SY5YQ94
B
Nuclei per well (x 1000)
0
2
4
6
***
***
***
Dox
CFZ
- - + + + + - +
